# Response to Comment on “Two types of asynchronous activity in networks of excitatory and inhibitory spiking neurons”

**DOI:** 10.1101/020354

**Authors:** Srdjan Ostojic

## Abstract

Networks of excitatory and inhibitory neurons form the basic computational units in the mammalian cortex. Within the dominant paradigm, neurons in such networks encode and process information by asynchronously emitting action potentials. In a recent publication, I argued that unstructured, sparsely connected networks of integrate-and-fire neurons display a transition between two qualitatively different types of asynchronous activity as the synaptic coupling is increased. A comment by Engelken et al (bioRxiv doi: 10.1101/017798) disputes this finding. Here I provide additional evidence for a transition between two qualitatively different types of asynchronous activity and address the criticism raised in the comment. The claims that the original paper is ”factually incorrect” and ”conceptually misleading” are unsubstantiated and inappropriate.

---

Excitatory-inhibitory networks of spiking neurons exhibit a well-studied asynchronous state in which individual neurons fire asynchronously and highly irregularly. In absence of temporal inputs, such irregular activity is conceptualized as being generated by a constant firing rate. Classical mean field theory provides a theoretical framework that formalizes this idea and relates quantitatively this firing rate to neural and network parameters.

The starting point of my recent paper [1] is the simple observation that the predictions of mean-field theory drastically break down when the coupling in the network is increased (Fig. P1a; the figures in the original paper are denoted by a number preceded by a ”P”). This salient fact has not been previously reported or explained. The central claim of the paper is that these strong deviations from classical mean-field theory can be interpreted as a signature of a transition from a state in which neurons fire at constant firing rates to a new type of asynchronous state in which the rates of different neurons *strongly fluctuate* both in time and across neurons. In a recent comment, Engelken et al claim to refute this interpretation by pointing out its limitations. They however do not provide an alternative explanation for the drastic breakdown of the classical mean-field description, which they do not dispute. Before replying to their specific criticisms, here I provide additional evidence for the central claim of the paper.

The interpretation proposed in the paper is based on an analogy between the spiking network and a network of rate units. This rate network can be understood as an extension of mean-field theory, in which every spiking neuron has its own firing rate. For low synaptic coupling, the activity in the rate model is constant and corresponds to the equilibrium fixed point of classical mean-field theory, i.e. classical asynchronous activity. As the coupling is increased beyond a critical value, an instability occurs and the rate network develops a state in which the firing rates strongly fluctuate both in time and across units. These fluctuations lead to strong deviations of the mean-firing rate away from the equilibrium value (Fig. P2). The crucial point is that these fluctuations are not weak fluctuations around a stable equilibrium: these are *strong* fluctuations in the sense that the they would diverge in absence of bounds on the activity. This implies that increasing the upper bound would increase fluctuations, which in turn would increase deviations of the mean-firing rate away from the equilibrium. The upper bound on firing rates can be easily modified in both rate and spiking models by varying the refractory period, and this simple manipulation leads to a qualitatively similar picture in the two models (Fig. 1). For low coupling strengths, the refractory period has no effect on mean-firing rates or fluctuations as instantaneous firing rates are well below saturation. Above a critical value of the coupling, as the refractory period is taken to zero both mean firing rates and fluctuations increase drastically (Fig. 1 **a,b**) and appear to diverge (Fig. 1 **c**). This behavior demonstrates directly (i.e. without reference to quantitative predictions of the mean-field theory) the existence of two qualitatively different types of asynchronous states, and validates the picture proposed in the paper. Again, the key difference between the two states is the *strength* of fluctuations.

**Figure 1:**
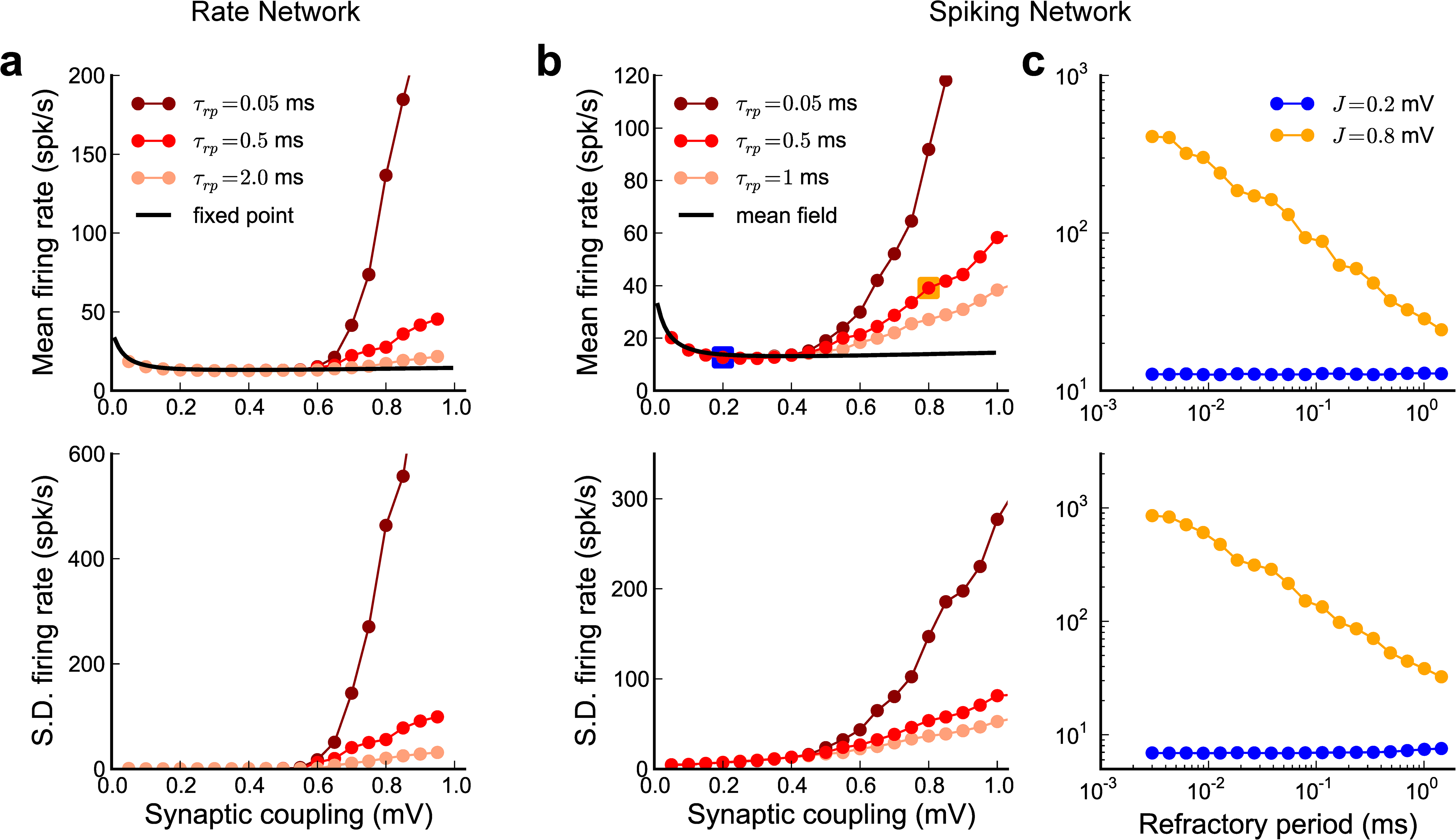
Effects of the upper bound on the activity in rate and spiking networks. **a**: Mean firing rate (top) and standard deviation of the firing rate (bottom) as the synaptic coupling is varied in the rate network, for three values of refractory period. **b**: Same as in **a**, for the spiking network. **c**: Mean firing rate (top) and standard deviation of the firing rate (bottom) as the refractory period is varied in the spiking network, for two values of synaptic coupling. Models and parameters as in the original paper, Figs. P1 and P2. In the spiking network, the standard deviation was computed on rates evaluated with 50-ms wide Gaussian filter (as in Fig. P3).

The analogy between the spiking network and the rate model is useful because it illustrates how network feedback leads to strongly fluctuating activity. This analogy is however a simplification that relies on a number of approximations, which are repeatedly pointed out in the original paper. The Comment by Engelken and colleagues nevertheless seems to choose a literal reading in which the spiking model would be exactly equivalent to the rate model. By pointing out differences between the two models, they claim to refute the existence of a transition between two different states.

The first point raised in the Comment et al are quantitative disagreements between the spiking and the rate models on the value of the critical coupling. Such quantitative disagreements are expected due to the approximate nature of the correspondence between the spiking and the rate model, and this is explicitly pointed out in the original article (Discussion, second paragraph of ”Generality of the findings”). The authors of the comment illustrate their point with a particular example of such a disagreement. Varying the refractory period for that specific parameter set nevertheless reveals that the transition is clearly present (Fig. 2 **a**). The predicted critical coupling is not accurate, but appears to be highly sensitive to the precise parameter values (Fig. 2 **b**).

**Figure 2:**
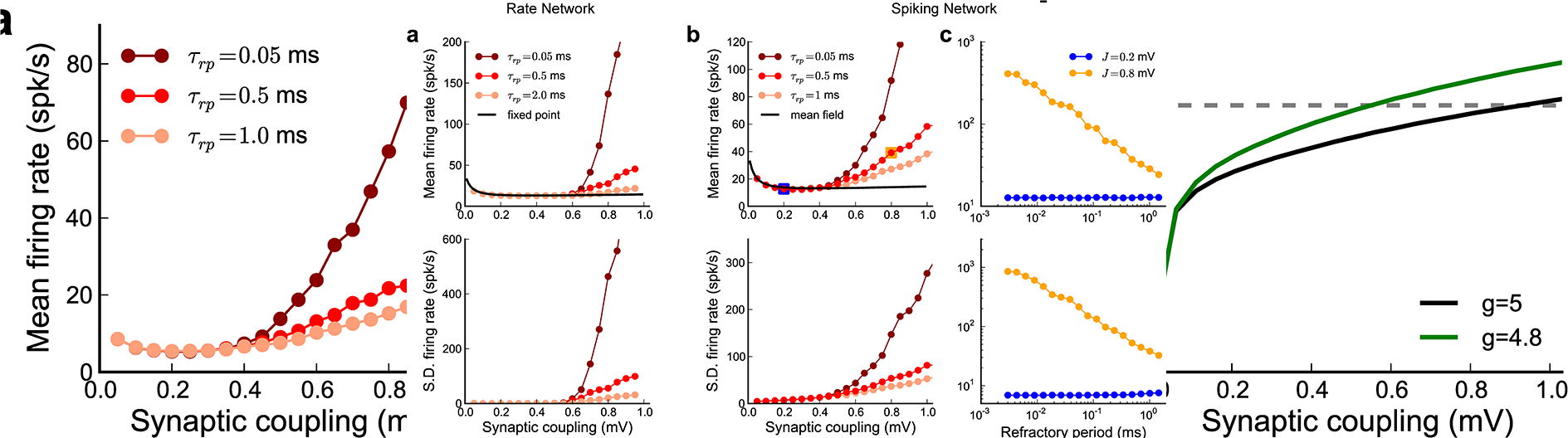
Results for the alternate parameter set used in the Comment. **a**: Mean firing rate (left) and standard deviation of the firing rate (right) as function of synaptic coupling in the spiking network, for three values of the refractory period. Following the Comment, the number of connections per neuron is *C* = 4000, and the size of the network is *N* = 40000, all other parameters as in Fig. P1. The standard deviation was computed on rates evaluated with 50-ms wide Gaussian filter (as in Fig. P3). **b**: Spectral radius as function of synaptic strength. For parameter values used in **a** (relative inhibition strength *g* = 5), the spectral radius predicts a critical coupling of about 1 mV. The critical coupling is however strongly sensitive to parameter values: if the relative inhibition strength is changed to *g* = 4.8, the critical coupling is down to 0.5 mV.

The second point raised in the Comment is that the transition apparently disappears when the synaptic delays are taken to zero. As pointed out in the article (Methods, below Eq.4), this is an artifact of instantaneous synaptic currents. A closer inspection of the activity for vanishing synaptic delays reveals that the network is highly synchronized (Fig. 3). Spikes emitted in synchrony reach other neurons during their refractory period (i.e. while the membrane potential dynamics are clamped to the reset potential), and since they are instantaneous have no effect at all, so that the neurons that fire in synchrony do not interact. The network is not at all in an asynchronous state, the analogy with a rate model is not valid and the whole picture proposed in the paper breaks down. As mentioned in the paper, this artifactual synchronization disappears as soon as delays are longer than the refractory period (Fig. 3 **b-c**), and the behavior then becomes independent of the precise value of the delay.

**Figure 3:**
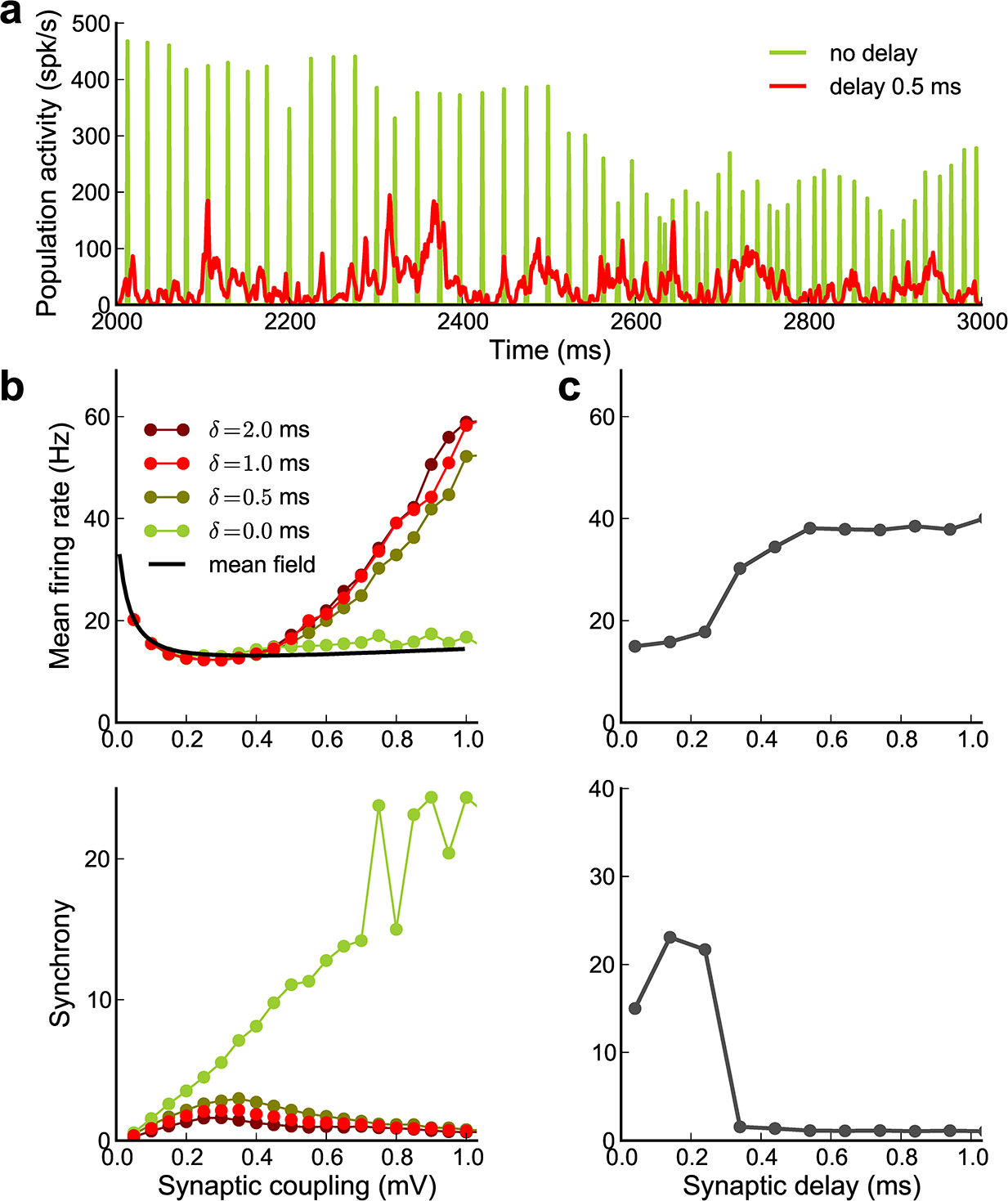
Effects of delays in the spiking network. **a**: Population activity (1 ms bins) for a synaptic coupling *J* = 0.8 mV, and two values of delay. **b**: Mean firing rate (top) and synchrony (bottom) as function of synaptic coupling, for four values of the synaptic delay. **c**: Mean firing rate (top) and synchrony (bottom) as function of synaptic delay, for a synaptic coupling of 0.8 mV. The amount of synchrony was quantified using the value of the population autocorrelation at zero lag (Eq. 21 in [1]).

The Comment then turns to the difference between the temporal aspects of the dynamics in the spiking and rate models. These differences are explicitly pointed out in the original paper (*“Spike-train autocorrelation functions in the LIF network display a short-timescale structure absent in the Poisson network”*). The absolute timescale in the rate model is totally arbitrary, and this is explicitly pointed out (Methods, below Eq.19). Similarly, the timescale of the instantaneous rates extracted by filtering from the spiking network is obviously arbitrary. The paper therefore does not make any statements on the precise *timescales* of the fluctuations, but on the increase in their *strength*. The authors of the comment instead normalize the amplitude of the fluctuations and focus on the precise shape of the auto-correlation functions. The precise shape of the auto-correlations is clearly different, and this is explicitly shown in the paper (Fig. P3 **c**).

The Comment next focuses on the close vicinity of the critical coupling. The authors correctly point out that the mechanism of the instability implies the existence of a diverging timescale at the critical coupling, and this should have been more clearly mentioned in the original article. In the rate model, this timescale manifests itself in the auto-correlation function: at the transition, the auto-correlation function changes from flat to very broad but of vanishing amplitude; above the transition its timescale quickly decreases as its amplitude increases (see Supplementary Figure in the Comment). In the spiking model, the auto-correlation is in contrast never flat, but nevertheless displays a qualitative change close to the transition: below the transition it is oscillatory, and around the transition a central peak emerges and increases in amplitude until the oscillations disappear (Fig. 4; the Comment shows only the auto-correlation functions well above the transition). A diverging time-scale is indeed not readily apparent in the auto-correlation in the spiking network. One possibility is that a slow component of vanishing magnitude is masked by the faster, oscillatory components. Another possibility is that the absence of diverging timescale in the auto-correlation is a genuine qualitative difference between the transitions in the spiking model and the proposed rate model. Note that other variants of the rate model exhibit the same type of transition, but without a diverging timescale in the auto-correlation function [3, 4] (see also Fig. 13 B in [5]). Moreover adding noise to the rate model eliminates the diverging timescale, as it leads to autocorrelation functions of finite amplitude and finite timescale.

**Figure 4:**
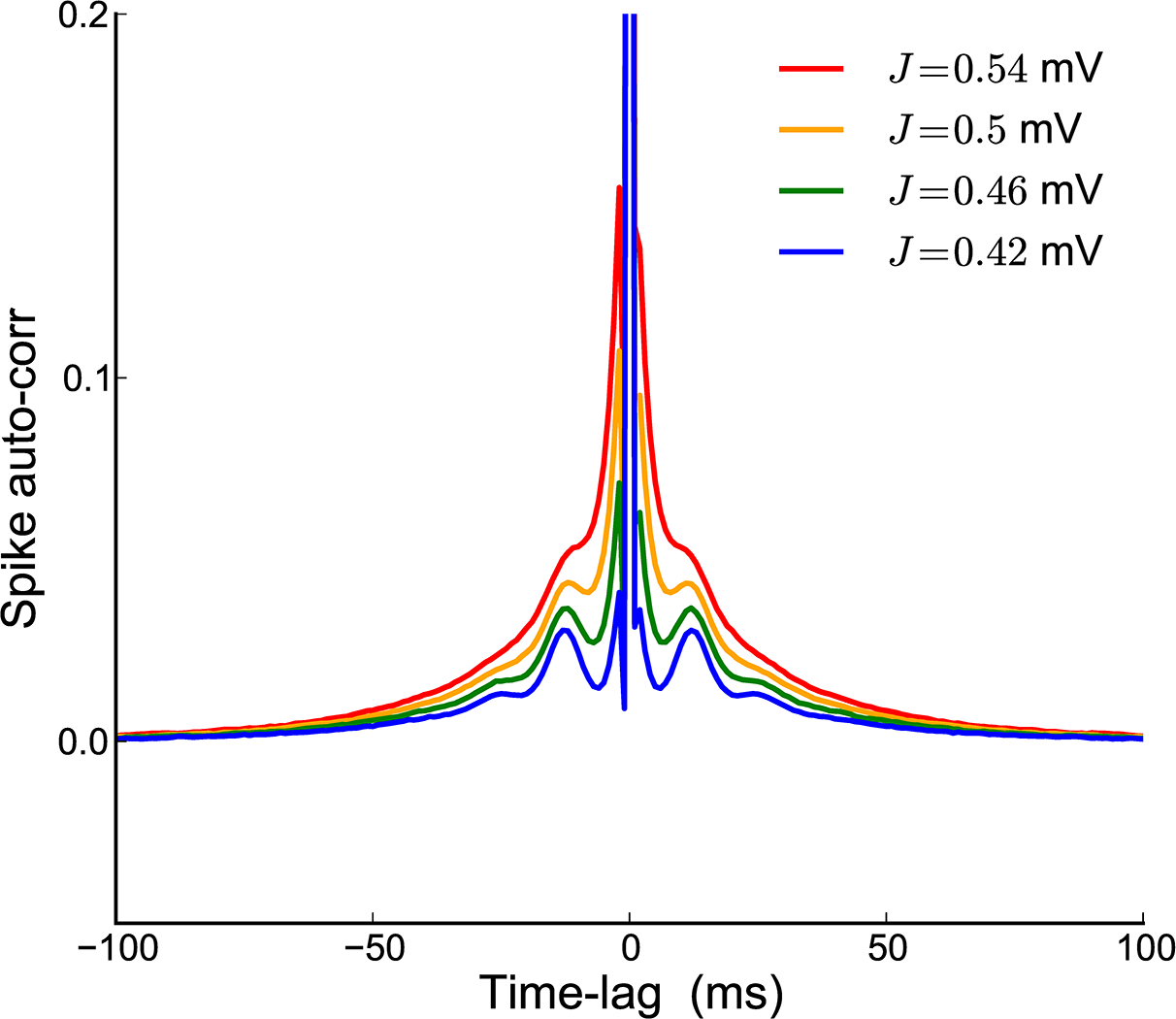
Auto-correlation functions in the spiking network close to the transition. All parameters as in Fig. P1. The auto-correlation functions are normalized as in Eq. 23 of the paper.

Finally the Comment examines the timescale of perturbations along the most unstable direction. The authors use a very quick pulse of 2 ms. If the amplitude of the pulse is vanishingly small, the rate model predicts an exponentially decaying response, the timescale of which diverges close to the instability. The authors argue that such a diverging time-scale is not seen in a network of leaky integrate-and-fire neurons. The response of leaky integrate-and-*v*fire neurons to a quick pulse is however never exponential, but contains a very fast component (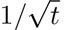 divergence in the impulse response at short times, see [6] p. 4). This response moreover strongly depends on the statistics of background fluctuations, and is expected to become even faster when background fluctuations are correlated in time [7], as is the case around the transition. A more detailed analysis is needed to understand how these various effects shape the network response.

The introduction of the Comment suggests that a transition to ”slow dynamics” is a central claim of the paper. Such a claim is never made in the original paper, which focuses on *strong* fluctuations, not *slow* fluctuations. The Comment also argues that very slow dynamics around the transition are essential for computations. As the Comment itself shows, the rate network exhibits very slow dynamics only if the coupling is very precisely tuned to the critical value (see supplementary figure in the Comment - the timescale decays over the range of 0.05 mV). The existing computational schemes [8, 9] do not rely on such fine-tuning as they do not exploit the diverging timescale, but rather the strong internal dynamics of the network [10], and this is the aspect that was stressed in the original paper. While very slow dynamics are certainly an interesting property, how networks (whether rate- or spike-based) robustly develop such dynamics is still an open question.

In summary, the results reported in the Comment by Engelken and colleagues do not challenge the main finding of the paper, the existence of two qualitatively different types of asynchronous activity. The Comment points out some limitations in the proposed picture, and in particular the behavior close to the transition point. These aspects certainly need to be better understood, but the Comment provides no additional or competing explanation for the phenomenon.

